# Neural signals to violations of abstract rules using speech-like stimuli

**DOI:** 10.1101/581355

**Authors:** Yamil Vidal, Perrine Brusini, Michela Bonfieni, Jacques Mehler, Tristan Bekinschtein

**Affiliations:** Cognitive Neuroscience Sector, International School for Advanced Studies (SISSA), Via Bonomea 265, 34136 Trieste, Italy; Institute of Psychology, Health and Society, University of Liverpool, Liverpool, United Kingdom; The University of Edinburgh, Edinburgh, EH8 9AD, United Kingdom; Department of Psychology, University of Cambridge, Cambridge, CB2 3EB, United Kingdom

## Abstract

As the evidence of predictive processes playing a role in a wide variety of cognitive domains increases, the brain as a predictive machine becomes a central idea in neuroscience. In auditory processing a considerable amount of progress has been made using variations of the Oddball design, but most of the existing work seems restricted to predictions based on physical features or conditional rules linking successive stimuli. To characterise the predictive capacity of the brain to abstract rules, we present here two experiments that use speech-like stimuli to overcome limitations and avoid common confounds. Pseudowords were presented in isolation, intermixed with infrequent deviants that contained unexpected phoneme sequences. As hypothesized, the occurrence of unexpected sequences of phonemes reliably elicited an early prediction error signal. These prediction error signals do not seemed to be modulated by attentional manipulations due to different task instructions, suggesting that the predictions are deployed even when the task at hand does not volitionally involve error detection. In contrast, the amount of syllables congruent with a standard pseudoword presented before the point of deviance exerted a strong modulation. Prediction error’s amplitude doubled when two congruent syllables were presented instead of one, despite keeping local transitional probabilities constant. This suggest that auditory predictions can be built integrating information beyond the immediate past. In sum, the results presented here further contribute to the understanding of the predictive capabilities of the human auditory system when facing complex stimuli and abstract rules.

**Significance Statement:** The generation of predictions seem to be a prevalent brain computation. In the case of auditory processing this information is intrinsically temporal. The study of auditory predictions has been largely circumscribed to unexpected physical stimuli features or rules connecting consecutive stimuli. In contrast, our everyday experience suggest that the human auditory system is capable of more sophisticated predictions. This becomes evident in the case of speech processing, where abstract rules with long range dependencies are universal. In this article, we present two electroencephalography experiments that use speech-like stimuli to explore the predictive capabilities of the human auditory system. The results presented here increase the understanding of the ability of our auditory system to implement predictions using information beyond the immediate past.

## Introduction

In recent years, the study of predictive processes has drawn increasing attention in neuroscience. In this context, Predictive Coding has emerged as a popular theory, which states that the brain constructs a hierarchy of predictions of incoming stimuli at multiple levels of processing (Bubic et al., 2010; Hobson and Friston, 2012; Friston, 2005, 2009, 2010). This proposal has received mounting empirical evidence (Wacongne et al., 2011; Phillips et al., 2015, 2016; Den Ouden et al., 2012).

A wealth of experiments in the study of Predictive Coding are variations of the OddBall design (Heilbron and Chait, 2018; Squires et al., 1975), where frequent acoustic stimuli establish predictable sequences, which are at times violated. Besides designs using tones, the use of speech-like stimuli offers a number of advantages. Within speech, abstract rules are ubiquitous, allowing to test abstract predictions that go beyond physical stimuli features and local transitional probabilities. These properties make speech processing an excellent testbed for the study of the brain’s signals to abstract rules establishment and its violations.

Speech perception requires the fast extraction of meaning from a complex auditory signal (Klein-schmidt and Jaeger, 2015; Boudewyn et al., 2015) and the generation of predictions might be an efficient solution to achieve fast and accurate comprehension (Kleinschmidt and Jaeger, 2015; Hauk, 2016). Although the proposal that predictive processes play a role in speech processing has been criticized (Norris et al., 2000; Van Petten and Luka, 2012; Huettig et al., 2015), evidence suggests that predictions are deployed at several speech processing levels (Lewis and Bastiaansen, 2015). At the syntactic level, listeners’ knowledge influence sentence parsing (Traxler, 2014; Wilson and Garnsey, 2009; Farmer et al., 2006; Baart and Samuel, 2015). Lexico-semantic processing can be facilitated by contextual predictability (Van Petten et al., 1999; Schuster et al., 2016).

Electroencephalography (EEG) studies have identified an event related potential (ERP) known as N400, whose amplitude is inversely correlated with the semantic predictability of words in context (Ku-tas and Hillyard, 1980; Van Petten et al., 1999; Brink et al., 2001; Kutas and Federmeier, 2000, 2011; Freunberger and Roehm, 2016; DeLong et al., 2005). EEG evidence has also shown that forthcoming phonemes can be predicted using syntactic (DeLong et al., 2005), semantic (Bendixen et al., 2014; Kashino, 2006; Groppe et al., 2010), phonological (Cornell et al., 2011; Hestvik and Durvasula, 2016; Schluter et al., 2016; Scharinger et al., 2016) and phonotactic information (Dehaene-Lambertz et al., 2000; Sun et al., 2015; Ylinen et al., 2016).

As the generation of predictions seem to be a prevalent brain computation (Friston, 2010, 2009), we propose that phonological predictions are generated during speech perception in the absence of semantic and syntactic information. To test this hypothesis, we performed 2 electroencephalography (EEG) experiments with an OddBall design. The use of speech stimuli allowed us to test for predictions based on an abstract rule that go beyond local transitional probabilities.

Pseudowords were presented in a context that did not contain syntactic or semantic information. We expected that the presentation of deviants, constructed using the same phonemes as standard pseudowords but in an unexpected sequence, would elicit an early prediction error signal like the Mismatch Negativity (MMN) (Näätänen et al., 2007; Winkler and Schröger, 2015; Friston, 2005; Chennu et al., 2013; Wacongne et al., 2011; Garrido et al., 2009). The presence of this prediction error signal would imply that listeners’ brains generate predictions about incoming phonemes within pseudoword.

We propose that abstract predictions are deployed regardless of the task at hand. To test this, experiments 1 and 2 differed with respect to the instructions given to the participants. While in experiment 1 participants were instructed to count the occurrence of deviants, in experiment 2 they were required to learn all pseudowords. We expected that an early prediction error signal would be present in both experiments, implying that predictions are deployed even if the task at hand does not require error detection and independent of the strategy of rule-learning.

Finally, in order to test if these predictions are constructed using information beyond local transitional probabilities, we tested whether the amplitude of prediction error would be modulated by the amount of phonemes presented before the point of deviance. We expected to find higher prediction error (higher amplitudes) when longer sequences of phonemes that are congruent with a standard pseudoword are presented. This modulation would not occur if predictions were made based solely on local transitional probabilities between phonemes.

Taken together, these experiments allowed us to study the predictive capabilities of the brain networks underlying the extraction of abstract rules.

## Materials and Methods

Stimuli set, unprocessed data and processing scripts can be found at https://osf.io/tuvy6/.

### Participants

Participants were self-reported right handed, Italian native speakers recruited from the city of Trieste with no auditory or language-related problems. Participants signed informed consent and received a monetary compensation of 15 e. Thirty participants (10 male, 20 female, mean aged 22.86 *±* 3.42 years) took part in experiment 1, and 29 participants (9 male and 20 female, mean aged 23.24 *±* 3.52 years) took part in experiment 2. After data preprocessing, participants contributing with less than 30 clean EEG trials per condition were excluded from analysis (1 participant excluded from each experiment). The remaining participants had sufficient trials to be included in a single subject statistical analysis and all contribute similarly to the group variance. Additionally, 1 participant was excluded from experiment 1 due to poor behavioural performance. Therefore, 28 participants (10 male, 18 female, mean age 23.25 *±* 3.23 years) from experiment 1 and 28 participants (8 male and 20 female, mean age 23.10 *±* 3.51 years) from experiment 2 were included in the final analyses.

### Stimuli

Six pseudowords divided in 3 sets of 2 pseudowords each were used as stimuli. We applied a series of constrains in the construction of our stimuli to ensure that the resulting pseudowords would resemble real Italian words. First we consulted the phonItalia lexical database (Goslin et al., 2014) to identify syllable candidates composed by 1 consonant followed by 1 vowel (i.e. 2 phonemes each). In order to exclude monosyllabic words and onomatopoeias, we removed syllables with a token frequency above the 70th percentile. Next, in order to keep syllables that could take any position within a word, we removed syllables with initial, middle or final position token frequencies either bellow the 20th percentile or above the 90th percentile. This selection procedure allowed us to identify 24 syllable candidates that are not monosyllabic words (in Italian) and have an even frequency distribution across positions within a word.

Using these syllable candidates, we constructed 2 trisyllabic pseudowords that contained no vowel or consonant repetitions. Additionally, no syllables were repeated between these 2 pseudowords. Hereafter, these pseudowords will be referred to as STD (i.e. Standard) pseudowords. Taking these STD pseudowords as a base, we constructed 2 different types of deviant pseudowords. The first deviant type, to which we will refer as XYY, consisted of the 1st syllable of a STD pseudoword and the 2nd and 3rd of the other STD pseudoword. The second type of deviant, to which we will refer as XXY, consisted of the 1st and 2nd syllable of a STD pseudoword, and the 3rd of the other STD pseudoword. Finally, 2 additional pseudowords with a XYX structure were constructed, only to be used as NEW pseudowords in a forced choice test at the end of experiment 2. None of these deviant pseudowords contained either consonant or vowel repetitions.

Audio file of these 2 STD pseudowords were generated using the MBROLA speech synthesizer (Dutoit et al., 1996) and the Italian female diphone database it4. Consonant and vowel durations were set to 150ms and 175ms respectively, hence, pseudowords duration was 975ms. Once the 2 STD pseudowords were produced, deviants were constructed by cross-splicing (i.e. cutting and replacing sound segments) the audio of the STD.

In natural speech, phonemes are co-articulated (i.e. the sound of each phoneme is influenced by the preceding and the forthcoming phoneme). Hence, using cross-splicing to generate the deviant pseudowords could result in sharp transitions that would sound unnatural. Because of this, we took measures to obtain a natural render for our stimuli (Steinberg et al., 2012). For the first and last syllable position, the vowels of both STD pseudowords had similar first and second formants. As one STD pseudoword had the vowel ‘o’ in the first syllable, the other STD pseudoword had the vowel ‘u’ at the same position. In the case of the third syllable, while one STD pseudoword used the vowel ‘i’, the other one used the vowel ‘e’. In the case of the second syllable, both STD pseudowords had ‘a’ as the vowel (Figure 1, A). For each syllable position, the consonants of both STD pseudowords had the same mode of articulation. Finally, the point of cutting was set close to zero amplitude. This measures had the effect of reducing the difference between both STD pseudowords at the points of syllable transitions so that when cross-spliced to construct the deviant pseudowords, these would not contain sharp transitions.

**Figure 1:**
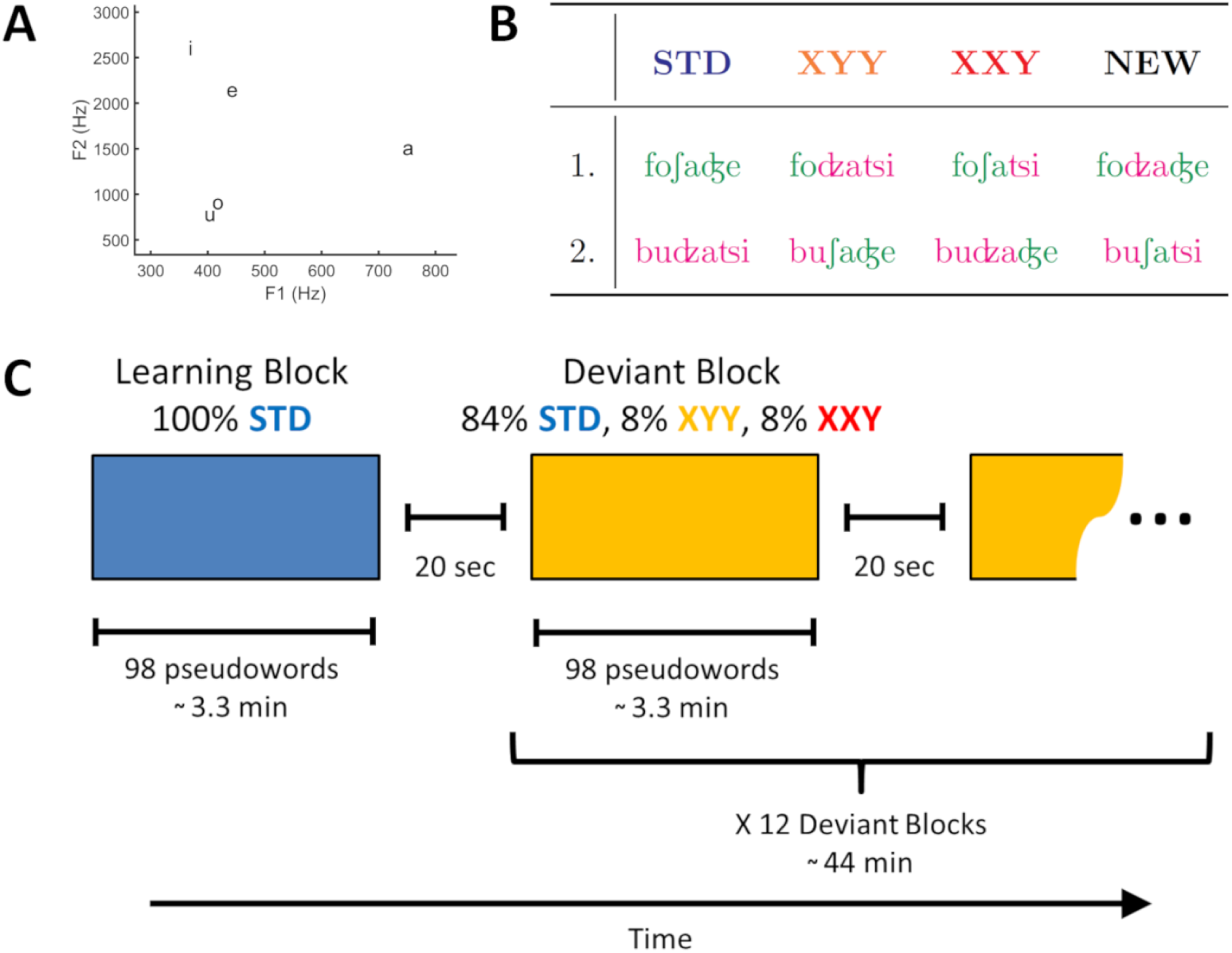
**A**: Scatter plot of 1st and 2nd formant of each vowel. **B:** Stimulus set in IPA notation. Deviant pseudowords were produced by cross-splicing the 2 STD pseudowords either at the end of the first syllable (XYY) or at the end of the second syllable (XXY). Two additional NEW pseudowords with a XYX structure were used only in a forced choice test at the end of experiment 2. **C:** In both experiments, stimuli were presented in 13 blocks separated by 20 seconds. Within each block, pseudowords were presented with an inter stimulus interval between 900 and 1300 ms. The first blocks consisted solely of STD pseudowords. Subsequent blocks were composed of 84% STD pseudowords 8% XYY deviant pseudowords and 8% XXY deviant pseudowords. Pseudoword order was pseudo-random. A minimum of 2 and a maximum of 4 STD pseudowords were presented between deviants and no deviants were presented more than 2 times consecutively.

The final set consisted of 2 STD, 2 XYY deviants, 2 XXY deviants and 2 NEW pseudowords (Figure 1, B). All pseudowords were checked by a native Italian speaker linguist to ensure that they sounded as plausible but not real Italian words.

While previous work in the literature has shown that the generation of predictions can serve word processing, phonemes in these experiments were either omitted (Bendixen et al., 2014), or replaced either by other phonemes (Cornell et al., 2013; Politzer-Ahles et al., 2016; Schluter et al., 2017) or by a non-linguistic sound (Kashino, 2006; Groppe et al., 2010). Because of this, changes in low level auditory features might have contributed to the recorded signals. In the case of our stimuli set, any difference in the EEG recording found between the STD condition and the deviant conditions could not be attributed to differences in instantaneous low level features. Instead, they could in principle only be attributed to the violation of the abstract rule learnt during the experiment (Paavilainen, 2013), according to which given a syllable *X*_*n*_, the next syllable of the word should be *X*_*n*+1_.

Note that in the case of the stimuli used here, the only feature that defined a pseudoword as deviant was that following the syllable *X*_*n*_, instead of the usual syllable *X*_*n*+1_, the syllable *Y*_*n*+1_ (which belongs to a different STD pseudoword) was presented. Additionally, as the overall frequency of presentation of all syllables used to construct the stimuli was the same, this design avoids a common confound between expectation and frequency of presentation (Heilbron and Chait, 2018).

### Experimental Design

Participants were requested to minimize movement throughout the experiment, except during breaks between blocks. No particular instructions were given with respect to when to blink, as eye blink artefacts can be removed using Independent Component Analysis (Delorme and Makeig, 2004; Chaumon et al., 2015).

Experiments followed an OddBall design, divided in 13 blocks with an average duration of 3.3 minutes each. During each block, a total of 98 pseudowords were presented, with an inter stimulus interval that varied between 900 and 1300 ms. During the first of such blocks, only STD pseudowords were presented. Subsequently, participants completed 12 blocks composed of 84% Standard pseudowords 8% XYY deviant pseudowords and 8% XXY deviant pseudowords. Within each block, pseudoword order was pseudo-random. A minimum of 2 and a maximum of 4 STD pseudowords were presented between deviants and no deviants were presented more than 2 times consecutively (Figure 1, C).

In experiment 1, participants were instructed to learn all made up “words” (i.e. pseudowords) in block one, and from block 2 onwards count the occurrence of “mistaken words” (i.e. deviant pseudowords) and write down the number of “mistaken” words during the pauses between blocks.

In experiment 2, participants were not informed about the presence of deviants and were simply instructed to learn all made up “words” (i.e. pseudowords). In order to ensure that the participant would pay attention during the experiment, they were informed that they would be subject to a test after the word learning task. After listening to the blocks of pseudowords, behavioural performance was assessed, by means of a forced choice test. On each trial, participants heard 2 pseudowords in sequence and were requested to choose the one that most likely was presented during the experiment. Participants completed 4 trials for each of 6 contrasts between conditions, for a total of 24 trials, presented in pseudorandom order (only 1 repetition of contrast type was allowed). The contrasts between conditions were “STD vs XYY”, “STD vs XXY”, “XYY vs XXY”, “STD vs NEW”, “XYY vs NEW” and “XXY vs NEW”. Participants reported their answers verbally and the experimenter entered them through keyboard. Order of presentation of pseudowords within trial was counterbalanced.

### Data acquisition setup

EEG data was collected using a 128 passive electrode system (Geodesic EEG System 300, Electrical Geodesics, Inc.) referenced to the vertex. EEG signal was bandpass filtered by hardware between 0.1 and 100 Hz, and digitalized at 250 Hz. Electrode impedance was kept below 100 kΩ(equivalent to 10kft standard amplifiers; Johnson et al., 2001). Participants were tested in a soundproof faraday cage while sitting on a chair in front of a LCD 19 inches monitor. Sound was delivered via a loudspeaker located behind the monitor, at a comfortable sound intensity of approximately 60 dB. Experiments were programmed in MATLAB (MathWorks, Inc., Natick, MA, USA, RRID: SCR_001622) using the Psychophysics Toolbox extensions (Brainard, 1997; Pelli, 1997) (RRID: SCR_002881). Pseudoword onset was marked on the EEG data by sending both a digital input signal (DIN) and a TCP/IP mark.

### EEG data pre-processing

EEG data preprocessing was performed in MATLAB using custom code and the EEGLAB toolbox (Delorme and Makeig, 2004) (RRID: SCR_007292). After being imported to EEGLAB, the data of each subject was band-pass filtered (0.1-30Hz). As the anti-aliasing filter of the EGI 300 Amp introduces a delay of 36ms, latencies of all events were corrected. The entire learning block, and the first 6 trials of each block, where excluded from analysis. Data was segmented into 1848ms long epochs starting 300ms before pseudoword onset. Bad channels were rejected using the 3 available methods of EEGLAB’s *pop_rejchan* function. *Kurtosis* threshold was set to 4σ, *Joint probability* threshold was set to 4σ, and *Abnormal spectra* was checked between 1 and 30 Hz, with a threshold of 3σ (Delorme and Makeig, 2004). Following this automatic cleaning, additional channels were rejected by visual inspection of continuous data and spectra. Independent Component Analysis (ICA) was use to remove eye blinks (Delorme and Makeig, 2004; Chaumon et al., 2015). Following, data was re-referenced to the average of all electrodes and baseline corrected using the 300ms before pseudoword onset. Next, we performed trial rejection by eliminating trials containing extreme values (*±* 200 *µ*V) and improbable trials (EEGLAB *pop_jointprob* 4σ for both *Single Channel* and *All Channels*). Finally, missing channels were interpolated (EEGLAB *pop_interp*, ‘spherical’).

Only after this cleaning procedure the data was divided into conditions. Given that STD pseudowords were presented far more frequently than deviant pseudowords, the datasets of each condition were pruned by randomly discarding trials to obtain exactly the same number of trials per condition. For example, if after trial rejection a participant had 763 STD trials, 76 XYY trials and 68 XXY trials, then 68 randomly picked trials per condition were kept and the rest were discarded. Participants contributing with less than 30 clean EEG trials per condition were excluded from analysis (one participant was excluded from each experiment applying this criterion). After this, the mean amount of trials per participant and condition were 70.18 *±* 16.57 (minimum = 35) for experiment 1 and 82.50 *±* 13.76 (minimum = 41) for experiment 2. For each condition, the mean of all trials of each subject was calculated and saved into a final dataset. The result of preprocessing was 1 dataset per condition, containing the mean of each subject.

Deviant conditions differed between each other with respect to the amount of syllables presented before the point of deviance. In order to render possible the comparison of the deviant conditions, we re-segmented the trials of both deviant conditions so that the points of deviance would be aligned. The resulting epochs had a length of 1224ms, starting 325ms before the point of deviance. Additionally, as the processing of a pseudoword has an intrinsic temporal dynamic, we eliminated this confounding factors by subtracting the activation elicited by the STD condition from each deviant condition.

### EEG Regions of Interest

Statistical analysis of EEG data was restricted to 2 predefined spatio-temporal Regions of Interest. The first one consisted on a Fronto-Central ROI comprised of 13 electrodes and spanned over a 325ms time window starting at the point of deviance of each deviant condition. With respect to word onset, this window spanned from 325ms to 650ms for the XYY condition, and from 650ms to 975ms for the XXY condition. This ROI coincided with the region were an early prediction error response like the MMN could be expected (Wacongne et al., 2012; Lecaignard et al., 2015; Bendixen et al., 2012; Duncan et al., 2009). The second region of interest consisted on a Parietal ROI composed of 21 electrodes and temporally extended from 200ms after the point of deviance of each deviant condition, to the end of the epoch. With respect to word onset, this window started at 525ms for the XYY condition, and at 850ms for the XXY condition. This ROI corresponded to the region were a P3b response would be expected (Comerchero and Polich, 1999; Polich, 2007; Duncan et al., 2009). As this component is strongly modulated by top-down attention (Sergent et al., 2005; Pegado et al., 2010; Dehaene and Changeux, 2011; Bekinschtein et al., 2009), it was used to test whether the attentional manipulation between experiments 1 and 2 was successful.

### Statistical Analysis

EEG group level contrast between conditions was performed utilizing a nonparametric clustering methods, introduced first by Bullmore et al. (1999) and implemented in the FieldTrip toolbox for EEG/MEG analysis (Oostenveld et al., 2011) (RRID: SCR_004849). This method offers a straightforward and intuitive solution to the Multiple Comparisons problem. It relies on the fact that EEG data has a spatio-temporal structure. A true effect should not be isolated but should instead spread over different electrodes and over time. Instead of assessing for differences between conditions in a point by point fashion, which would lead to a very big number of comparisons, this method groups together adjacent spatio-temporal points.

The procedure is as follows. For every point in time and space, the EEG signal of 2 conditions is statistically compared. In our case, we used a nonparametric permutation *t* test for this step. The *t* values of adjacent spatio-temporal points with *p* values *<* 0.05 are clustered together and a cluster-level statistic is calculated by summing the *t* values within a cluster. Once these candidate clusters have been defined, their probability of occurrence under the null hypothesis of no difference between conditions is assessed using a nonparametric permutation test. In this test, conditions are shuffled and cluster-level *t* values are calculated as before. This step is repeated 5000 times and on each iteration the most extreme cluster-level *t* value is retained. This allows to construct an histogram of expected cluster-level *t* values under the null hypothesis of no difference between the conditions. Cluster level *p* values are calculated as the proportion of expected *t* values under the null hypothesis that are more extreme than the observed *t* value. For further details see Maris and Oostenveld (2007).

Additionally, in order to corroborate results found at the group level were robust and not driven by outliers, we performed a test at the participant level. For each individual participant, the mean amplitude over the time of the detected group level cluster was calculated, and the conditions of interest were submitted to a paired *t* test in order to obtain a *t* value. Next, the *t* values from all participants were converted to 1 if they show a difference between conditions in the same direction as the group lever cluster or 0 if otherwise. A one-tailed binomial test was performed on these transformed *t* values, with equal or lower likelihood as null hypothesis. The logic of this analysis is that if an effect is true at the group level, then the majority of participants should show a difference between conditions in the same direction. Note that the test used is one-tailed because the hypothesis to test is directional.

All effect sizes reported are Hedges’ *g* (Hedges, 1981; Lakens, 2013), which is less biased than Cohen’s *d*, as it applies a correction for small sample sizes. Effect sizes were calculated using the Measures of Effect Size Toolbox (Hentschke and Stüttgen, 2011). Additional statistical analysis were performed using JASP version 0.8.6 (Bayesian analyses; JASPteam, 2017) and RStudio version 1.1.456 (Linear mixed effects models; RStudioTeam, 2016) (RRID: SCR_000432).

## Results

Given that deviant conditions differed in the time point at which a pseudoword could be identified as a deviant (325ms and 650ms from pseudoword onset for XYY and XXY conditions respectively), instead of defining time zero as onset of stimulus presentation, we will use the time point of deviance of each conditions as such. In other words, all times reported are with respect to the point of deviance. Furthermore, comparisons across deviants and experiment were performed on the difference wave between STD and deviant, and with all trials re-segmented to align the point of deviance, as described in Materials and Methods.

### Behavioural results

In experiment 1, participants were requested to count the occurrence of mistaken words (i.e deviant pseudowords) on each block. On average, participants reported 15.22 (out of 16 presented) deviant pseudowords per block (*σ* = 2.56). For each participant, we checked the number of blocks with a deviant count further than 2*σ* from the mean. While most of the participants reported a deviant count within these limits for all the blocks, 3 participants had 1 block with a lower count, and 1 participant had all 12 blocks outside this limit. This participant reported a mean of only 3.58 deviants per block, therefore, was excluded from the analysis. After excluding this participant and 1 other participant that contributed with less than 30 clean EEG trials per condition, the mean number of deviants reported per block increases to 15.62 (*σ* = 1.41). This performance is close to ceiling (16).

Note that the method of asking participants to mentally count the occurrence of deviants does not allow us to determine with certainty neither the occurrence of false alarms, nor the detection rate for each deviant condition. Despite this, given that the mean count of deviant was close to the actual number of deviants presented, we can conclude that in experiment 1, participants were able to perform the task with high accuracy for both deviant conditions.

Contrary to experiment 1, during experiment 2 participants were not aware of the presence of deviant pseudowords. Despite this, at the end of the experiment, they were requested to perform a force choice test in which each stimuli condition was contrasted against the others and against new pseudowords not presented during the blocks. The mean preference in each contrast was calculated for each participant and a one sample *t* test was performed at the group level to test against the null hypothesis of no difference from chance (i.e. 50%). Results were corrected for multiple comparisons using the Bonferroni-Holm method.

Participants preferred STD pseudowords over both deviant pseudoword types. They choose STD pseudowords over XYY deviants on 67.24% of the trials (*t*_(28)_ = 3.57, *p* = 0.0051, *g* = 0.66 [0.25, 1.06]) and over XXY deviants on 69.82% of the trials (*t*_(28)_ = 4.07, *p* = 0.0017, *g* = 0.75 [0.33, 1.16]). When both deviant types were contrasted, participants preferred XYY over XXY deviants on 62.06% of the trials, but this preference was not reliable (*t*_(28)_ = −2.31, *p* = 0.056, *g* = 0.43 [0.04, 0.80]).

Next, we contrasted the pseudowords used in the experiment against NEW pseudowords that were not previously presented. Participants preferred STD pseudowords over NEW pseudowords on 85.34% of the trials (*t*_(28)_ = 10.39, *p* = 2.46*×*10^*-*10^, *g* = 1.92 [1.30, 2.54]) and XXY deviants over new pseudowords on 64.65% of the trials (*t*_(28)_ = 2.99, *p* = 0.0169, *g* = 0.55 [0.16, 0.94]). XYY deviants on the contrary, could not be distinguished from NEW pseudowords as they were preferred on only 55.17% of the trials (*t*_(28)_ = 1.03, *p* = 0.3117, *g* = 0.19 [-0.17, 0.55]).

These behavioral results allowed us to corroborate that participants paid attention during the blocks of pseudowords. They also indicate that in experiment 2, despite the fact that the instructions provided did not explicitly distinguish between standard and deviant pseudowords, participants displayed a preference for STD pseudowords over both deviant pseudoword types. Even though both deviant types had the same probability of occurrence, while XXY deviants could be distinguished from NEW pseudowords, XYY could not. Taken together these behavioral results suggest that participants were sensitive to the frequency of occurrence of the different pseudowords.

### EEG evidence of abstract rule extraction via phonological predictions

In order to test whether phonological predictions are deployed during speech perception in the absence of semantic and syntactic information, we used clustering (see Materials and Methods) to compared each deviant condition against the STD condition, focusing the analysis on the Fronto-Central ROI, where the presentation of a deviant pseudoword was expected to elicit an early prediction error signal.

In experiment 1, XYY deviants elicited such response, peaking in amplitude at 155ms (*t*_(27)_ = −89.77, *p* = 0.0004, *g* = −0.90 [-1.35, −0.44]), followed by a positive deflection with peak amplitude at 227ms (*t*_(27)_ = 36.81, *p* = 0.0208, *g* = 0.57 [0.09, 1.05]) (Figure 2, A). XXY deviant also elicited a prediction error response with peak amplitude at 170ms (*t*_(27)_ = −125.20, *p* = 0.0002, *g* = −0.91 [-1.35, −0.47]), followed by a positive deflection with peak amplitude at 246ms (*t*_(27)_ = 57.88, *p* = 0.0126, *g* = 0.60 [0.21, 1.00]) (Figure 2, B).

**Figure 2:**
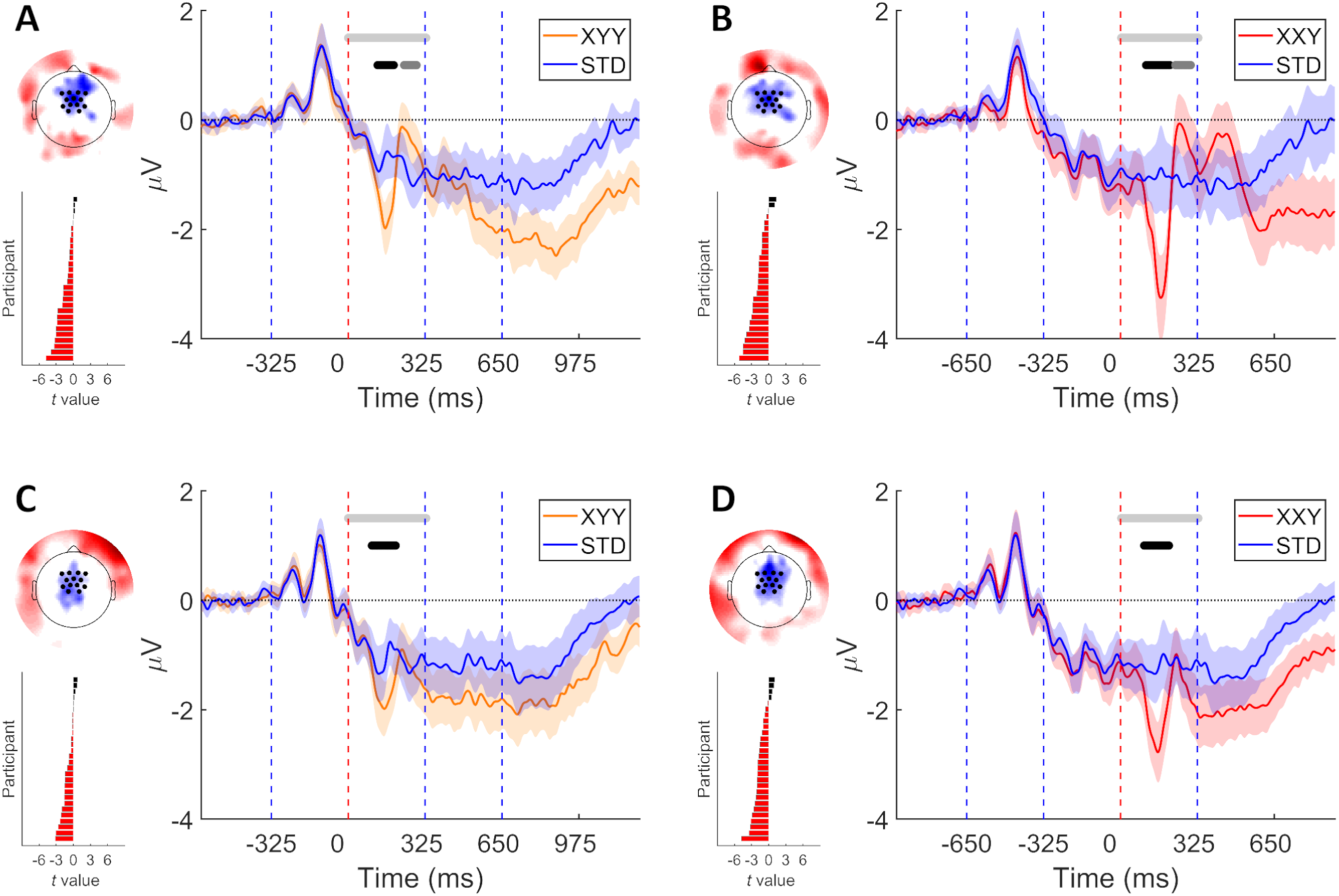
Early prediction error elicited by both deviant types in experiments 1 (**A** and **B**) and 2 (**C** and **D**). On each panel: Right, grand average over fronto-central ROI. Vertical dashed lines indicate syllable boundaries. Time zero indicates the point at which deviance occur. Shaded areas denote 95% CI. Horizontal light grey line delimits time window of analysis. Middle grey horizontal line indicates *p <* 0.05 (cluster corrected). Black horizontal line indicates *p <* 0.01 (cluster corrected). Left top, topography of the difference wave, mean over the time of the negative cluster. Left bottom, individual participants’ t values calculated over mean cluster time.

The results of experiment 1 show that the presentation of a deviants pseudoword, composed by an unexpected sequence of syllables, elicited prediction error signals. Since in experiment 1 participants were instructed to count “mistaken” (i.e. deviant) pseudowords, we sought to replicate these results under conditions more akin to natural speech perception. Experiment 2, while using the same stimuli and OddBall design of experiment 1, differed with respect to the instructions given to the participants. In experiment 2 participants were asked to learn all pseudowords, without informing them of the presence of deviants.

Once more our analysis of the Fronto-Central ROI revealed that both deviant types evoked a prediction error signal. XYY deviants elicited a response peaking in amplitude at 151ms (*t*_(27)_ = −100.79, *p* = 0.0004, *g* = −0.54 [-0.85, −0.23]) (Figure 2, C). In the case of XXY deviants, peak amplitude was reached at 158ms (*t*_(27)_ = −138.13, *p* = 0.0006, *g* = −0.78 [-1.14, −0.41]) (Figure 2, D).

Results at the group level were corroborated by performing a test participant by participant, as described in the Methods section. This analysis showed that in both experiments and for both deviant conditions, the majority of the participants displayed a difference between conditions in the direction congruent with the tested hypothesis (Experiment 1: XYY deviant, 24/28 85.71% *p* = 9*×*10^*-*5^; XXY deviant, 26/28 92.86% *p* = 1.52*×*10^*-*6^. Experiment 2: XYY deviant, 22/28 78.57% *p* = 0.00186; XXY deviant, 24/28 85.71% *p* = 9e-05).

Taken together, the results of experiments 1 and 2 show that the presentation of deviants composed by an unexpected sequence of syllables trigger an early prediction error signal. The presence of this error signal indicates that a prediction about the forthcoming syllables had been made, even when the context didn’t contain any syntactic or semantic information.

### Neural signals to violations of abstract rules under different instructions

In order to test whether predictions are deployed regardless of the task at hand, experiments 1 and 2 used the same stimuli and design, but differed in the instructions given to the participants. While in experiment 1 participants were requested to count the occurrence of deviants, in experiment 2 they were not informed about the presence of deviants and were instead requested to learn all pseudowords. Despite this difference, as we reported at the beginning of this section, the presentation of deviant pseudowords elicited an early prediction error signal in both experiments.

To confirm that the change in instructions successfully induced a different attention allocation between experiments, we analysed the signal recorded at the parietal ROI. If the attentional manipulation was successful, the presentation of a deviant pseudoword should elicit a P3b response only in experiment 1, where deviant detection was relevant for the task at hand (Bekinschtein et al., 2009).

In experiment 1, our analysis of the parietal ROI revealed that both deviant types elicited the expected P3b response. In the case of the XYY deviant, P3b response started at 251ms and reached 50% of its area under the curve at 743ms (*t*_(27)_ = 1625.05, *p* = 0.0002, *g* = 1.77 [1.03, 2.51]) (Figure 3, A). In turn, the P3b response elicited by the XXY deviant started at 262ms and reached 50% of its area under the curve at 578ms (*t*_(27)_ = 1149.20, *p* = 0.0002, *g* = 1.97 [1.17, 2.76]) (Figure 3, B). Furthermore, the amplitude of the P3b component was modulated by deviant type. XXY deviants elicited a higher amplitude P3b response than XYY deviants (*t*_(27)_ = 225.97, *p* = 0.0018, *g* = 0.47 [0.13, 0.80]) (Figure 4, C). This comparison was performed on the difference wave between STD and each deviant condition, with the point of deviance temporally aligned.

**Figure 3:**
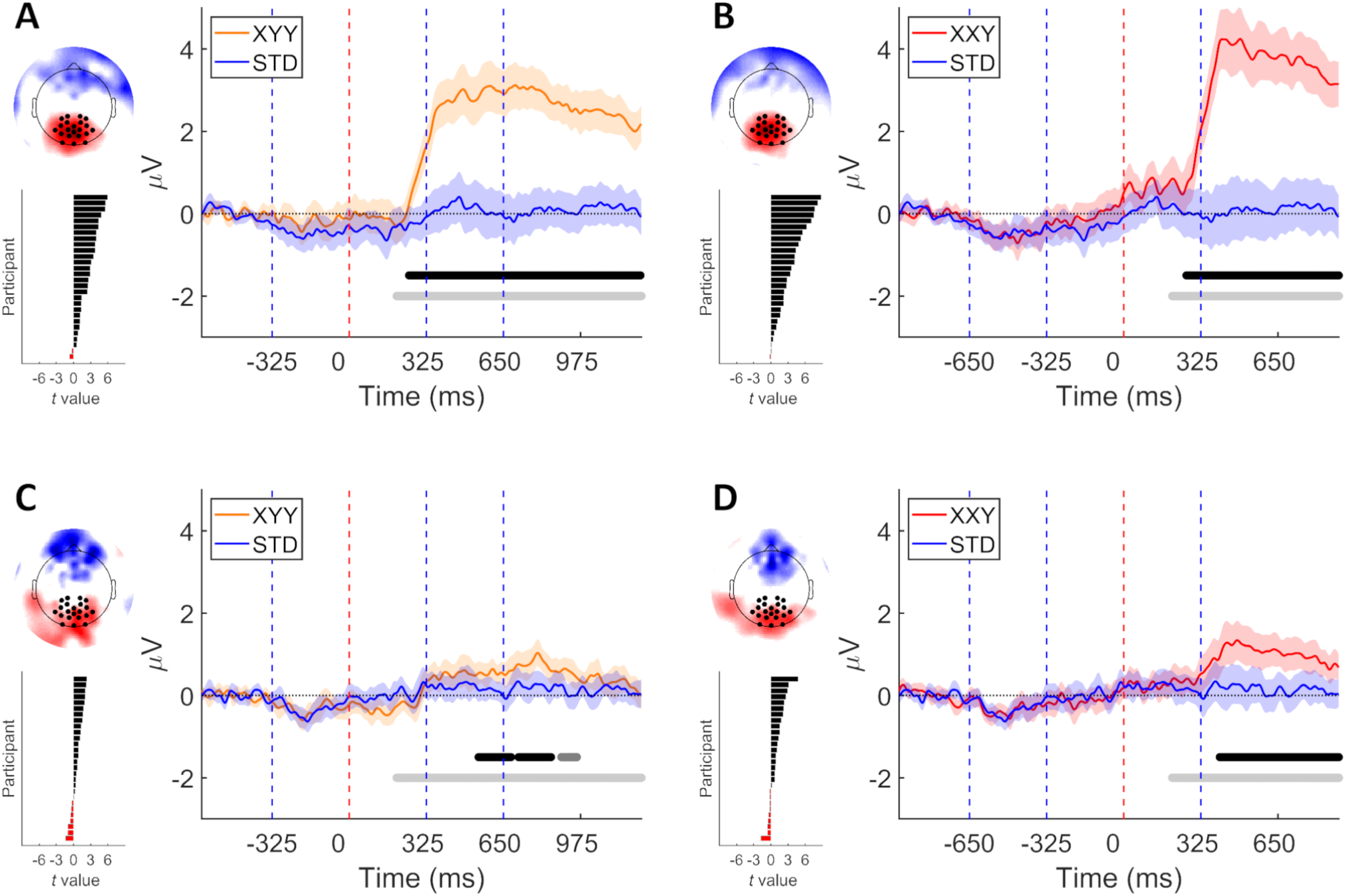
**A** P3b response was elicited by both deviant types in experiments 1 (**A** and **B**), but not detected in experiment 2 (**C** and **D**). On each panel: Right, grand average over parietal ROI. Vertical dashed lines indicate syllable boundaries. Time zero indicates the point at which deviance occur. Shaded areas denote 95% CI. Horizontal light grey line delimits time window of analysis. Middle grey horizontal line indicates *p <* 0.05 (cluster corrected). Black horizontal line indicates *p <* 0.01 (cluster corrected). Left top, topography of the difference wave, mean over the time of the positive cluster. Left bottom, individual participants’ t values calculated over mean cluster time.

**Figure 4:**
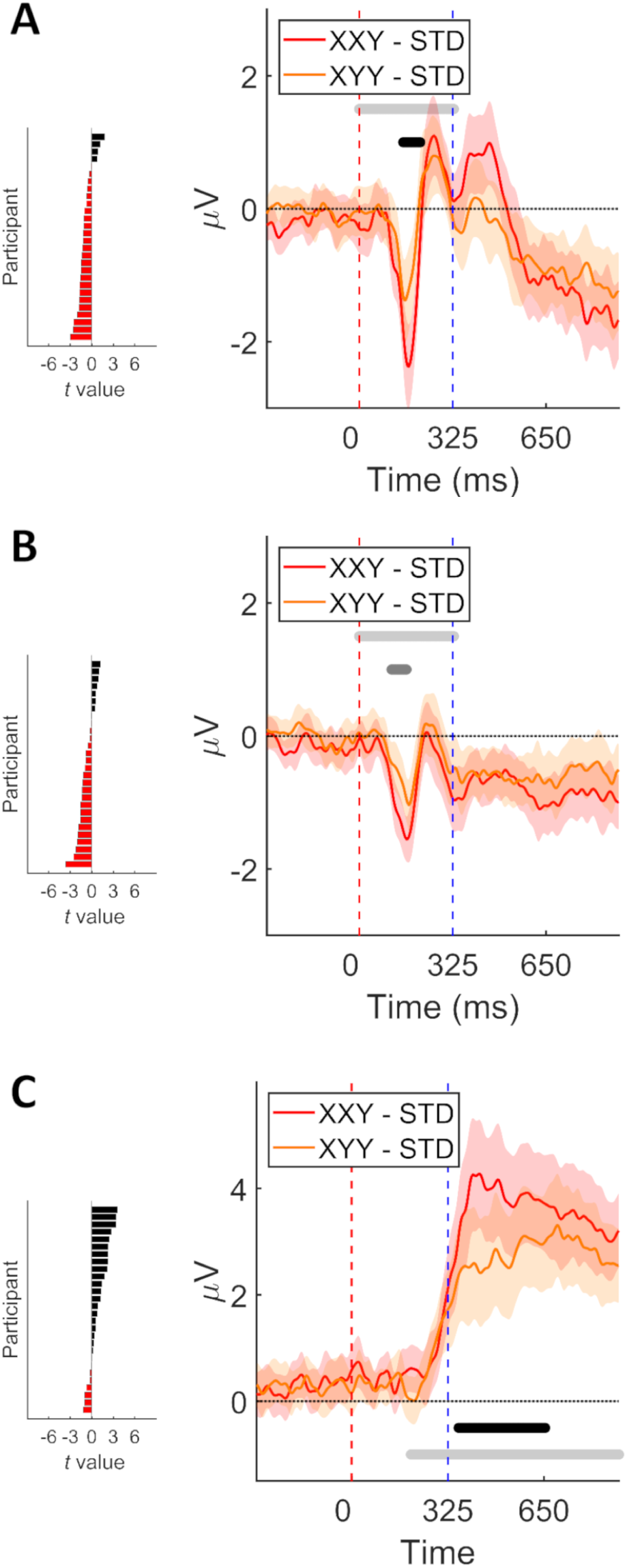
Comparison of signals elicited by each deviant type (Difference waves, deviant minus STD). On each panel: Right, grand average over fronto-central ROI (**A** and **B**), or parietal ROI (**C**). Trials were re-segmented and locked to the point of deviance, indicated by time zero. Shaded areas denote 95% CI. Horizontal light grey line delimits time window of interest. Middle grey horizontal line indicates *p <* 0.05 (cluster corrected). Black horizontal line demarks *p <* 0.01. Early prediction error signals detected in experiments 1 (**A**) and 2 (**B**). P3b detected in experiment 1 (**C**). Left, individual participants’ t values calculated over mean cluster time.

While in experiment 1 both deviant conditions elicited a clear P3b response, in experiment 2, only positivites of lower amplitude were detected. For the XYY deviant, clustering analysis detected a series of 3 consecutive positive clusters, the first of which started at 543ms (*t*_(27)_ = 99.39, *p* = 0.0092, *g* = 0.70 [0.15, 1.26]; *t*_(27)_ = 94.46, *p* = 0.0098, *g* = 0.60 [0.11, 1.09]; *t*_(27)_ = 44.64, *p* = 0.0362, *g* = 0.65 [0.11, 1.20]). In the case of the XXY deviant condition, a single positive cluster was found, starting at 402ms and reaching 50% of its area under the curve at 394ms (*t*_(27)_ = 385.57, *p* = 0.0014, *g* = 0.82 [0.21, 1.43]).

Next, to further confirm that the attentional manipulation between experiments was successful, we contrasted the recorded signals across experiments using clustering analysis. We expected to find higher amplitudes in experiment 1, due to the presence of the P3b elicited by the deviants. We were able to confirm this for both deviants (XYY: *t*_(54)_ = 875.00, *p* = 0.0002, *g* = 1.41 [0.80, 1.99]; XXY: *t*_(54)_ = 734.07, *p* = 0.0002, *g* = 1.26 [0.66, 1.82]). Analyses were performed on the difference between STD and deviant conditions. These results confirm that the top-down attention paid to deviants was indeed different between experiments.

Having confirmed that the attentional manipulation between experiments was successful, and considering that regardless of this, an early prediction error signal was registered in both experiments, we decided to test if the prediction error signals recorded across experiments where indeed equivalent. As our hypothesis stated that there would be no difference in prediction error amplitude across experiments (i.e. a null hypothesis), a Bayesian independent samples *t* test (Bayes Factor, Rouder et al., 2009) was used for these comparisons. This test measures the relative evidence between the null and alternative hypothesis, allowing to assess evidence in favour of the null (Leppink et al., 2017). Tests were performed using a Cauchy prior with scale value of *r* = 1.

We compared the amplitude of the early prediction error signals registered over the Fronto-Central ROI, elicited by each deviant condition across experiments, by taking the mean amplitude in a 44ms time window (equal to the duration of the shortest cluster) centred at the peak of the detected negativity. For both deviant types, Bayes Factor showed only anecdotal evidence in favour of no difference between experiments (XYY deviants: BF_01_ = 2.48, *g* = 0.32 [-0.20, 0.85]; XXY deviants: BF_01_ = 1.14, *g* = 0.48 [-0.06, 1.01]). Analyses were performed on the difference between STD and deviant conditions.

Taken together, these results suggest that even if the task at hand does not explicitly imply deviance detection, phonological predictions are proactively deployed. However it should be noted that the results with respect to the modulation of early prediction error by top down attention are inconclusive.

### Predictions beyond local transitional probabilities

The prediction error signals described above could reflect violations of predictions based on local transitional probabilities, or alternatively these predictions could be constructed by considering information in a longer cognitive time window. To shed light on this issue, we contrasted conditions where deviance occurred at different time points within a pseudoword. The logic behind this comparison is that if predictions are built not solely on the basis of local transitional probabilities, an increase in the number of syllables presented before the point of deviance would elicit higher amplitude prediction error signals. In XXY the second syllable lends further evidence that the pseudoword is about to be completed, but then this prediction is violated in the last syllable, while in XYY the prediction is broken earlier.

In both experiments the early prediction error signal elicited by XXY deviants had a bigger amplitude than the signal elicited by XYY deviants (Experiment 1: *t*_(27)_ = −53.95, *p* = 0.0094, *g* = −0.64 [-1.06, - 0.23]. Experiment 2: *t*_(27)_ = −38.02, *p* = 0.0204, *g* = −0.62 [-1.12, −0.12]; Figure 4, A and B). This suggests that prediction strength can be modulated by the amount of preceding syllables that are congruent with a STD pseudoword. Once more, we corroborated these results by performing a test participant by participant, as described in the Methods section. This analysis showed that in both experiments, the majority of the participants displayed higher amplitude prediction error signals for the XXY deviant (Experiment 1: 24/28 85.71% *p* = 9e-05; Experiment 2: 21/28 75.00% *p* = 0.00627).

It remained possible that small discrepancies in the number of STD trials presented before the deviants of each condition might be in part driving these effects. To rule out this possible confound, we fitted linear mixed effects models (using the *lme4* package in RStudio, Bates et al., 2015; RStudioTeam, 2016) to predict single trial prediction error amplitude using deviant type and amount of preceding STD trials (STD count) as fixed factors, and including participant as random factor (*PE* ∼ *Dev* + *STD*_*count* +(1 + *Dev*|*Participant*); *R*^2^ Experiment 1 = 0.0124, *R*^2^ Experiment 2 = 0.0116). An effect of deviant type was found in both experiments (Experiment 1: β = −0.94, *t*_(3929)_ = −3.705, *p* = 0.00021; Experiment 2: β = −0.67, *t*_(4619)_ = −3.530, *p* = 0.00042). In contrast, no effects of STD count were found (Experiment 1: β = −0.007, *t*_(3929)_ = −0.092, *p* = 0.92; Experiment 2: β = 0.059, *t*_(4619)_ = 1.172, *p* = 0.24). These results rule out the possibility that a substantial part of the difference in prediction error amplitude between deviant conditions would be driven by a difference in mean STD count preceding the deviants.

## Discussion

As we argued in the Introduction, the experimental designs typically used to study prediction in auditory processing share a number of limitations. The majority of the experimental designs used are variations of the Oddball paradigm (Heilbron and Chait, 2018). In most of these experimental designs, what defines a particular stimulus as deviant is the disruption of an established physical feature such as pitch, duration, intensity, side of stimulation or the presence of a gap (Näätänen et al., 2007). This limitation applies to the classical Oddball paradigm, optimum-1 (Näätänen et al., 2004), omission (Yabe et al., 1997) and roving-standard (Garrido et al., 2008) designs.

While these designs define standard and deviant stimuli on the basis of their physical features, other designs explore the sensitivity of the predictive system to higher order regularities or abstract rules that define the relationship between successive stimuli. For example Paavilainen et al. (2007) presented to their participants sequences of sinusoidal tone pips for which the duration varied randomly between short (50 ms) and long (150 ms). Importantly, the duration of each tone predicted the pitch of the next one, which could be either low (1000Hz) or high (1500 Hz). The authors found that the violation of this arbitrary abstract rule, linking duration of a tone with pitch of the next, elicited an early error signal (MMN response). Other examples of paradigms that test for prediction of higher order regularities are the unexpected repetition (Wacongne et al., 2012) and repetition vs expectation (Todorovic and de Lange, 2012) designs (for a review of abstract rule designs see Paavilainen, 2013).

Abstract rule designs have given support to Predictive Coding by showing that putative early prediction error signals, like the MMN response, cannot be fully explained by simple adaptation to standard stimuli (and lack of adaptation to deviant stimuli). But in all the designs mentioned above, the rules used established relationships only between consecutive stimuli. Therefore, these experimental designs only allow to study the sensitivity of the predictive system to local transitional probabilities.

To the best of our knowledge, there are only two paradigms that allow to test violations of an abstract rule beyond local transitional probabilities. In the Local/Global paradigm (Bekinschtein et al., 2009), tones are presented in groups of five. This allows to establish regularities both locally (transitional probabilities between tones within groups) and globally (between groups change, only tractable over a time range of seconds). In the RAND-REG designs (Barascud et al., 2016), tones are presented in succession at multiple possible pitches, switching between randomness and regular patterns. In these experiments, the detection of a regular pattern requires to consider several consecutive tones (1 full cycle plus 4 tones according to an ideal observer model). While the Local/Global and RAND-REG designs allow to study predictions that integrate information beyond adjacent stimuli, these designs use tone stimuli that are far less complex than naturally occurring sounds.

As evidence suggests that the generation of predictions might be one of the strategies that the speech processing system uses to parse the speech signal (Kleinschmidt and Jaeger, 2015; Hauk, 2016; Hickok, 2012; Norris et al., 2015; Boudewyn et al., 2015), and given that abstract rules and long range dependencies are ubiquitous in language, one way to overcome the limitations of the experimental designs described above is to use speech-like stimuli.

In the context of speech processing, it has been shown that listeners tend to hallucinate the presence of phonemes replaced by tones. The strength of this illusion depend on how much the preceding context is informative about the missing phoneme (Kashino, 2006; Groppe et al., 2010). Similarly, when a phoneme is omitted from a word (Bendixen et al., 2014), this can elicit a Mismatch Negativity (MMN) (Näätänen et al., 2007), which is a marker of violation of expectations (Friston, 2005; Winkler and Schröger, 2015), but only if the context in which the phoneme omission occurs contains semantic information that makes the omitted phoneme predictable. Phoneme replacements can also elicit a MMN response when the replacement violates a phonotactic rule of the language of the listener (Dehaene-Lambertz et al., 2000; Sun et al., 2015; Ylinen et al., 2016). Furthermore, and particularly framed in the context of Predictive Coding, it has been shown that the amplitude of the MMN response elicited by phoneme replacement is modulated by the availability of phonological evidence (i.e. degree of feature specification) of the preceding standard words before the presentation of a deviant (Scharinger et al., 2012a,b, 2016).

The studies described in the previous paragraph have provided compelling evidence of the role that predictions play in speech processing, but besides using speech as a complex auditory stimuli, they incorporate in their designs other linguistic factors such as syntax, semantic information and phonotactics. We proposed that phonological prediction might be generated within words, even in the absence of these additional sources of information. To test this, we performed 2 EEG OddBall experiments in which only phonological information was available to generate phonological predictions. Importantly, the deviant pseudowords used in these experiments were constructed by cross-splicing standard pseudowords. Therefore, each phoneme in a deviant pseudoword was acoustically identical to a phoneme in a standard pseudoword. The only feature that defined a pseudoword as deviant, was that following the syllable *X*_*n*_, instead of the usual syllable *X*_*n*+1_, the syllable *Y*_*n*+1_, which belongs to a different pseudoword, was presented. In this way, the ERP responses registered in these experiments could not be elicited by low frequency of occurrence of a given sound, or a change in instantaneous low level auditory features, but by the violation of an abstract rule (Paavilainen, 2013). As the stimuli did not contain consecutive phoneme repetitions, the registered responses cannot be explained by stimulus specific adaptation. Additionally, this stimuli design avoids a common confound between repetition and expectation (Todorovic and de Lange, 2012; Heilbron and Chait, 2018).

In both of the experiments presented here, the occurrence of an unexpected sequence of phonemes, reliably elicited an early prediction error signal, compatible with a MMN response (Näätänen, 2000; Näätänen et al., 2007). This event related potential is a well-established prediction error signal that can be interpreted as the result of comparing a prediction with the actual bottom-up input (Chennu et al., 2013; Wacongne et al., 2011; Garrido et al., 2009; Friston, 2005; Paavilainen, 2013; Winkler and Czigler, 2012). The presence of this early prediction error signal, elicited by the presentation of an unexpected sequence of phonemes, can be considered as evidence that a prediction about the forthcoming phonemes had been made.

Experiments 1 and 2 differed in the instructions given to the participants. While in experiment 1 participants were instructed to count the occurrence of “mistaken words” (i.e. deviants), in experiment 2 they were not informed about the occurrence of deviants and were simply instructed to learn all the pseudowords. This aimed to induce in experiment 2, an attentional state that resembles more closely the one held during natural speech processing.

To confirm the effects of this attentional manipulation, we tested for the presence of a P3b component in both experiments. While clustering analysis detected clear P3b components in experiment 1, only smaller positivities were detected in experiment 2. This suggest that participants noted the difference in frequency of occurrence between STD and deviant pseudowords, even when they were not instructed to detect deviants. In line with this, the behavioural results from experiment 2 show that participants preferred STD pseudowords over both deviant pseudoword types.

Despite this, when contrasting the signals recorded between experiments, we could verify that the amplitude in the P3b time window was roughly four times higher in experiment 1. As the P3b component is an index of to top-down attention (Chennu and Bekinschtein, 2012; Sergent et al., 2005; Bekinschtein et al., 2009; Dehaene and Changeux, 2011; Strauss et al., 2015; Faugeras et al., 2011), this difference indicates that the degree to which top-down attention was deployed was different between experiments.

In spite of the difference in instructions and in concomitant top-down attention between experiments, unexpected sequence of phonemes reliably elicited an early prediction error signal. This suggests that phonological predictions can be deployed, even if the task at hand does not require detecting abnormalities in the speech stream. Given that the results of our Bayesian analysis comparing amplitude of prediction error across experiments were inconclusive, the modulatory role that top-down attention might exert on these predictions remains an open question. As the attention allocation held by the participants during experiment 2 resembles closely the one use for natural speech processing, these results imply that the language comprehension system proactively anticipates incoming phonemes within individual words.

One way in which these phonological predictions could be implemented is by extracting the local transitional probabilities between adjacent syllables (Endress and Mehler, 2009; Koelsch, 2016). Our data indicates that this is unlikely, as we found that the amplitude of prediction error signals was modulated by the amount of syllables presented before the point of deviance. When 2 congruent syllables were presented before the point of deviance (XXY) the amplitudes were higher than when only 1 congruent syllable was presented (XYY). As the local transitional probabilities between *X*_1_ and *X*_2_ were the same as between *X*_2_ and *X*_3_ (.92), this increase in amplitude indicates that the information used to generate predictions was not restricted to consecutive syllables. Instead prediction strength was modulated by integrating information from several past phonemes.

It has been shown that the number of phonological features differing between standard and deviants can modulate the amplitude of the MMN response (Cornell et al., 2013; Scharinger et al., 2016; Schluter et al., 2017). Taking this into account, the difference in prediction error amplitude between deviant conditions may be captured by this feature. Taking the position of Mioni (1993) and Kramer (2009), who propose that in the case of Italian, affricates does not constitute a separate class of manner of articulation, the phonological features that change from STD to deviant in our stimuli set are the following. Syllables in the 2nd position (XYY deviant) differ in their consonant voicing, place of articulation and manner of articulation. Syllables in the 3rd position (XXY deviant) differ in their consonant voicing and place of articulation, and in their vowel height (Mioni, 1993; Kramer, 2009; Paoli, 2016). While it should be noted that whether all these phonological features have a neural representation is on itself an open debate (Hestvik and Durvasula, 2016; Politzer-Ahles et al., 2016; Schluter et al., 2016, 2017), in the case of our stimuli set, the number of phonological features that change for each deviant condition is the same.

Finally, when the point of deviance is reached, more time has elapsed from pseudoword onset in the case of XXY deviants, compared to XYY deviants. This difference in time from pseudoword onset could contribute to the difference in MMN amplitude, but we find this improbable. Behavioral gating experiments (Tyler, 1984) and MEG experiments (Brodbeck et al., 2018) have shown that between 50 to 100ms from word onset are enough to generate a prediction regarding the initial phoneme of a word. In the case of XYY deviants, the point of deviance is reached 325ms after pseudoword onset, which is more than 3 times the suggested minimum time for prediction generation. Therefore, the difference in elapsed time before deviance between conditions is unlikely to contribute to the observed difference in prediction error amplitude.

One tentative interpretation for the difference in prediction error amplitude between deviant conditions is that, as language processing is characterized by extensive communication across representational levels (Kuperberg and Jaeger, 2016; Davis and Johnsrude, 2007), a lexical level of processing could be involved. Specifically, when a phoneme of a word is perceived, this could be used to pre-activate that word’s lexical representation, with consecutive phonemes reinforcing the prediction of congruent words.

Taken together our results suggest that even when no higher level linguistic information such as syntax and semantics is present, the human auditory system can use phonological information from several past phonemes to generate predictions about forthcoming phonemes. In the experiments presented here, participants were exposed to new pseudowords that were learned in a period of minutes. This implies a formidable capacity of the auditory system to learn sequences of phonemes composing new words and generate predictions within those words. This capacity might play a fundamental role in the difficult task of mapping a complex, variable and noisy signal as speech into meaning. Moreover, the experiments presented here use stimuli and abstract rules more complex and ecologically valid that the ones routinely used in the study of auditory prediction, allowing to show that the auditory system can proactively generate predictions.

## Notes

https://osf.io/tuvy6/

## References

Baart, M. and Samuel, A. G. (2015). Early processing of auditory lexical predictions revealed by ERPs. Neuroscience Letters, 585:98–102.

Barascud, N., Pearce, M. T., Griffiths, T. D., Friston, K. J., and Chait, M. (2016). Brain responses in humans reveal ideal observer-like sensitivity to complex acoustic patterns. Proceedings of the National Academy of Sciences, 113(5):E616–E625.

Bates, D., Mächler, M., Bolker, B., and Walker, S. (2015). Fitting Linear Mixed-Effects Models Using lme4. Journal of Statistical Software, 67(1):1–48.

Bekinschtein, T. A., Dehaene, S., Rohaut, B., Tadel, F., Cohen, L., and Naccache, L. (2009). Neural signature of the conscious processing of auditory regularities. Proceedings of the National Academy of Sciences of the United States of America, 106(5):1672–7.

Bendixen, A., SanMiguel, I., and Schröger, E. (2012). Early electrophysiological indicators for predictive processing in audition: a review. International journal of psychophysiology : official journal of the International Organization of Psychophysiology, 83(2):120–31.

Bendixen, A., Scharinger, M., Strauß, A., and Obleser, J. (2014). Prediction in the service of comprehension: Modulated early brain responses to omitted speech segments. Cortex, 53:9–26.

Boudewyn, M. A., Corlett, P. R., Friston, K., Brown, M., and Kuperberg, G. R. (2015). A Hierarchical Generative Framework of Language Processing: Linking Language Perception, Interpretation, and Production Abnormalities in Schizophrenia. Front. Hum. Neurosci, 9(November):1–23.

Brainard, D. (1997). The Psychophysics Toolbox. Spatial Vision, 10(4):433–436.

Brink, V. D., Brown, C. M., and Hagoort, P. (2001). Electrophysiological Evidence for Early Contextual Influences during Spoken-Word Recognition : N200 Versus N400 Effects. Journal of Cognitive Neuroscience, 13(7):967–985.

Brodbeck, C., Hong, L. E., and Simon, J. Z. (2018). Rapid Transformation from Auditory to Linguistic Representations of Continuous Speech. Current Biology, 28(24):3976–3983.e5.

Bubic, A., von Cramon, D. Y., and Schubotz, R. I. (2010). Prediction, cognition and the brain. Frontiers in human neuroscience, 4(March):25.

Bullmore, E. T., Suckling, J., Overmeyer, S., Rabe-Hesketh, S., Taylor, E., and Brammer, M. J. (1999). Global, voxel, and cluster tests, by theory and permutation, for a difference between two groups of structural MR images of the brain. IEEE transactions on medical imaging, 18(1):32–42.

Chaumon, M., Bishop, D. V., and Busch, N. A. (2015). A practical guide to the selection of independent components of the electroencephalogram for artifact correction. Journal of Neuroscience Methods, 250:47–63.

Chennu, S. and Bekinschtein, T. a. (2012). Arousal Modulates Auditory Attention and Awareness: Insights from Sleep, Sedation, and Disorders of Consciousness. Frontiers in Psychology, 3(MAR):65.

Chennu, S., Noreika, V., Gueorguiev, D., Blenkmann, A., Kochen, S., Ibáñez, A., Owen, A. M., and Bekinschtein, T. A. (2013). Expectation and attention in hierarchical auditory prediction. The Journal of neuroscience : the official journal of the Society for Neuroscience, 33(27):11194–205.

Comerchero, M. D. and Polich, J. (1999). P3a and P3b from typical auditory and visual stimuli. Clinical Neurophysiology, 110(1):24–30.

Cornell, S. A., Lahiri, A., and Eulitz, C. (2011). “What you encode is not necessarily what you store": Evidence for sparse feature representations from mismatch negativity. Brain Research, 1394:79–89.

Cornell, S. A., Lahiri, A., and Eulitz, C. (2013). Inequality across consonantal contrasts in speech perception: Evidence from mismatch negativity. Journal of Experimental Psychology: Human Perception and Performance, 39(3):757–772.

Davis, M. H. and Johnsrude, I. S. (2007). Hearing speech sounds: Top-down influences on the interface between audition and speech perception. Hearing Research, 229(1-2):132–147.

Dehaene, S. and Changeux, J. P. (2011). Experimental and Theoretical Approaches to Conscious Processing. Neuron, 70(2):200–227.

Dehaene-Lambertz, G., Dupoux, E., and Gout, A. (2000). Electrophysiological correlates of phonological processing: A cross-linguistic study. Journal of Cognitive Neuroscience, 12(4):635–647.

DeLong, K. a., Urbach, T. P., and Kutas, M. (2005). Probabilistic word pre-activation during language comprehension inferred from electrical brain activity. Nature neuroscience, 8(8):1117–1121.

Delorme, A. and Makeig, S. (2004). EEGLAB: an open source toolbox for analysis of single-trial EEG dynamics including independent component analysis. Journal of neuroscience methods, 134(1):9–21.

Den Ouden, H. E. M., Kok, P., and de Lange, F. P. (2012). How prediction errors shape perception, attention, and motivation. Frontiers in Psychology, 3(DEC):1–12.

Duncan, C. C., Barry, R. J., Connolly, J. F., Fischer, C., Michie, P. T., Näätänen, R., Polich, J., Reinvang, I., and Van Petten, C. (2009). Event-related potentials in clinical research: Guidelines for eliciting, recording, and quantifying mismatch negativity, P300, and N400. Clinical Neurophysiology, 120(11):1883–1908.

Dutoit, T., Pagel, V., Pierret, N., Bataille, F., and Vrecken, O. V. D. (1996). The MBROLA project: towards a set of high quality speech\nsynthesizers free of use for non commercial purposes. Proceeding of Fourth International Conference on Spoken Language Processing. ICSLP ‘96, 3:2–5.

Endress, A. D. and Mehler, J. (2009). The surprising power of statistical learning: When fragment knowledge leads to false memories of unheard words. Journal of Memory and Language, 60(3):351–367.

Farmer, T. a., Christiansen, M. H., and Monaghan, P. (2006). Phonological typicality influences on-line sentence comprehension. Proceedings of the National Academy of Sciences of the United States of America, 103(32):12203–12208.

Faugeras, F., Rohaut, B., Weiss, N., Bekinschtein, T. a., Galanaud, D., Puybasset, L., Bolgert, F., Sergent, C., Cohen, L., Dehaene, S., and Naccache, L. (2011). Probing consciousness with event-related potentials in the vegetative state. Neurology, 77(3):264–8.

Freunberger, D. and Roehm, D. (2016). Semantic prediction in language comprehension: evidence from brain potentials. Language, Cognition and Neuroscience, 3798(July):1–13.

Friston, K. (2005). A theory of cortical responses. Philosophical transactions of the Royal Society of London. Series B, Biological sciences, 360(1456):815–36.

Friston, K. (2009). The free-energy principle: a rough guide to the brain? Trends in Cognitive Sciences, 13(7):293–301.

Friston, K. (2010). The free-energy principle: a unified brain theory? Nature reviews. Neuroscience, 11(2):127–138.

Garrido, M. I., Friston, K. J., Kiebel, S. J., Stephan, K. E., Baldeweg, T., and Kilner, J. M. (2008). The functional anatomy of the MMN: A DCM study of the roving paradigm. NeuroImage, 42(2):936–944.

Garrido, M. I., Kilner, J. M., Stephan, K. E., and Friston, K. J. (2009). The mismatch negativity: a review of underlying mechanisms. Clinical neurophysiology : official journal of the International Federation of Clinical Neurophysiology, 120(3):453–63.

Goslin, J., Galluzzi, C., and Romani, C. (2014). PhonItalia: a phonological lexicon for Italian. Behavior research methods, 46(3):872–86.

Groppe, D. M., Choi, M., Huang, T., Schilz, J., Topkins, B., Urbach, T. P., and Kutas, M. (2010). The phonemic restoration effect reveals pre-N400 effect of supportive sentence context in speech perception. Brain Research, 1361:54–66.

Hauk, O. (2016). Preface to special issue “prediction in language comprehension and production”. Language, Cognition and Neuroscience, 31(1):1–3.

Hedges, L. V. (1981). Distribution Theory for Glass’s Estimator of Effect size and Related Estimators. Journal of Educational and Behavioral Statistics, 6(2):107–128.

Heilbron, M. and Chait, M. (2018). Great Expectations: Is there Evidence for Predictive Coding in Auditory Cortex? Neuroscience, 389:54–73.

Hentschke, H. and Stüttgen, M. C. (2011). Computation of measures of effect size for neuroscience data sets. European Journal of Neuroscience, 34(12):1887–1894.

Hestvik, A. and Durvasula, K. (2016). Neurobiological evidence for voicing underspecification in English. Brain and Language, 152:28–43.

Hickok, G. (2012). The cortical organization of speech processing: feedback control and predictive coding the context of a dual-stream model. Journal of communication disorders, 45(6):393–402.

Hobson, J. A. and Friston, K. J. (2012). Waking and dreaming consciousness: neurobiological and functional considerations. Progress in neurobiology, 98(1):82–98.

Huettig, F., Mani, N., and Huettig, F. (2015). Is prediction necessary to understand language? Probably not. Language, Cognition, and Neuroscience, 3798(January):1–26.

JASPteam (2017). JASP (Version 0.8) [Computer software].

Johnson, M. H., Haan, M. D., Oliver, A., and Smith, W. (2001). Recording and Analyzing High-Density Event-Related Potentials With Infants Using the Geodesic Sensor Net. Developmental neuropsychology, 19(3):295–323.

Kashino, M. (2006). Phonemic restoration: The brain creates missing speech sounds. Acoustical Science and Technology, 27(6):318–321.

Kleinschmidt, D. and Jaeger, T. F. (2015). Robust speech perception: Recognize the familiar, generalize to the similar, and adapt to the novel. Psychological Review, 122(2):148–203.

Koelsch, S. (2016). Under the hood of statistical learning: A statistical MMN reflects the magnitude of transitional probabilities in auditory sequences. Scientific reports, 6(August 2015):1–11.

Kramer, M. (2009). The Phonology of Italian. Oxford University Press, Oxford, England.

Kuperberg, G. R. and Jaeger, T. F. (2016). What do we mean by prediction in language comprehension? Language Cognition & Neuroscience, 31(1):32–59.

Kutas, M. and Federmeier, K. D. (2011). Thirty Years and Counting: Finding Meaning in the N400 Component of the Event-Related Brain Potential (ERP). Annual Review of Psychology, 62(1):621–647.

Kutas, M. and Federmeier, K. K. D. (2000). Electrophysiology reveals semantic memory use in language comprehension. Trends in Cognitive Sciences, 4(12):463–470.

Kutas, M. and Hillyard, S. (1980). Reading senseless sentences: brain potentials reflect semantic incongruity. Science, 207(4427):203–205.

Lakens, D. (2013). Calculating and reporting effect sizes to facilitate cumulative science: A practical primer for t-tests and ANOVAs. Frontiers in Psychology, 4(NOV):1–12.

Lecaignard, F., Bertrand, O., Gimenez, G., Mattout, J., and Caclin, A. (2015). Implicit learning of predictable sound sequences modulates human brain responses at different levels of the auditory hierarchy. Frontiers in Human Neuroscience, 9(September):505.

Leppink, J., O’Sullivan, P., and Winston, K. (2017). Evidence against vs. in favour of a null hypothesis. Perspectives on Medical Education, 6(2):115–118.

Lewis, A. G. and Bastiaansen, M. (2015). A predictive coding framework for rapid neural dynamics during sentence-level language comprehension. Cortex, 68:155–168.

Maris, E. and Oostenveld, R. (2007). Nonparametric statistical testing of EEG- and MEG-data. Journal of Neuroscience Methods, 164(1):177–190.

Mioni, A. (1993). Fonetica e fonologia. In Sobrero, A., editor, Introduzione all’italiano contemporaneo, Le strutture, pages 101–139. Laterza, Rome.

Näätänen, R. (2000). Mismatch negativity (MMN): Perspectives for application. International Journal of Psychophysiology, 37(1):3–10.

Näätänen, R., Paavilainen, P., Rinne, T., and Alho, K. (2007). The mismatch negativity (MMN) in basic research of central auditory processing: a review. Clinical neurophysiology : official journal of the International Federation of Clinical Neurophysiology, 118(12):2544–2590.

Näätänen, R., Pakarinen, S., Rinne, T., and Takegata, R. (2004). The mismatch negativity (MMN): Towards the optimal paradigm. Clinical Neurophysiology, 115(1):140–144.

Norris, D., McQueen, J. M., and Cutler, A. (2000). Merging information in speech recognition: Feedback is never necessary. Behavioral and Brain Sciences, 23(3):299–325.

Norris, D., McQueen, J. M., and Cutler, A. (2015). Prediction, Bayesian inference and feedback in speech recognition. Language, Cognition and Neuroscience, 3798(December 2015):1–15.

Oostenveld, R., Fries, P., Maris, E., and Schoffelen, J.-M. (2011). FieldTrip: Open source software for advanced analysis of MEG, EEG, and invasive electrophysiological data. Computational intelligence and neuroscience, 2011:156869.

Paavilainen, P. (2013). The mismatch-negativity (MMN) component of the auditory event-related potential to violations of abstract regularities: A review. International Journal of Psychophysiology, 88(2):109–123.

Paavilainen, P., Arajärvi, P., and Takegata, R. (2007). Preattentive detection of nonsalient contingencies between auditory features. NeuroReport, 18(2):159–163.

Paoli, S. (2016). A short guide to Italian Phonetics and Phonology for students of Italian Paper V. Oxford University.

Pegado, F., Bekinschtein, T., Chausson, N., Dehaene, S., Cohen, L., and Naccache, L. (2010). Probing the lifetimes of auditory novelty detection processes. Neuropsychologia, 48(10):3145–54.

Pelli, D. (1997). The VideoToolbox software for visual psychophysics: transforming numbers into movies. Spatial Vision, 10(4):437–442.

Phillips, H. N., Blenkmann, A., Hughes, L. E., Bekinschtein, T. A., and Rowe, J. B. (2015). Hierarchical Organization of Frontotemporal Networks for the Prediction of Stimuli across Multiple Dimensions. Journal of Neuroscience, 35(25):9255–9264.

Phillips, H. N., Blenkmann, A., Hughes, L. E., Kochen, S., Bekinschtein, T. A., Cam, C. A., and Rowe, J. B. (2016). Convergent evidence for hierarchical prediction networks from human electrocorticography and magnetoencephalography. Cortex, 82:192–205.

Polich, J. (2007). Updating P300: an integrative theory of P3a and P3b. Clinical neurophysiology : official journal of the International Federation of Clinical Neurophysiology, 118(10):2128–48.

Politzer-Ahles, S., Schluter, K., Wu, K., and Almeida, D. (2016). Asymmetries in the perception of Mandarin tones: Evidence from mismatch negativity. Journal of Experimental Psychology: Human Perception and Performance, 42(10):1547–1570.

Rouder, J. N., Speckman, P. L., Sun, D., Morey, R. D., and Iverson, G. (2009). Bayesian t tests for accepting and rejecting the null hypothesis. Psychonomic Bulletin and Review, 16(2):225–237.

RStudioTeam (2016). RStudio: Integrated Development Environment for R.

Scharinger, M., Bendixen, A., Trujillo-Barreto, N. J., and Obleser, J. (2012a). A sparse neural code for some speech sounds but not for others. PLoS ONE, 7(7).

Scharinger, M., Monahan, P. J., and Idsardi, W. J. (2012b). Asymmetries in the Processing of Vowel Height. Journal of Speech, Language, and Hearing Research, 55(3):903–918.

Scharinger, M., Monahan, P. J., and Idsardi, W. J. (2016). Linguistic category structure influences early auditory processing: Converging evidence from mismatch responses and cortical oscillations. NeuroImage, 128:293–301.

Schluter, K., Politzer-Ahles, S., and Almeida, D. (2016). No place for /h/: An ERP investigation of English fricative place features. Language, Cognition and Neuroscience, 31(6):728–740.

Schluter, K. T., Politzer-Ahles, S., Al Kaabi, M., and Almeida, D. (2017). Laryngeal features are phonetically abstract: Mismatch negativity evidence from Arabic, English, and Russian. Frontiers in Psychology, 8(MAY):1–19.

Schuster, S., Hawelka, S., Hutzler, F., Kronbichler, M., and Richlan, F. (2016). Words in Context: The Effects of Length, Frequency, and Predictability on Brain Responses During Natural Reading. Cerebral Cortex, 26 (10)(June):1–16.

Sergent, C., Baillet, S., and Dehaene, S. (2005). Timing of the brain events underlying access to consciousness during the attentional blink. Nature neuroscience, 8(10):1391–400.

Squires, N. K., Squires, K. C., and Hillyard, S. A. (1975). Two varieties of long-latency positive waves evoked by unpredictable auditory stimuli in man. Electroencephalography and Clinical Neurophysiology, 38:387–401.

Steinberg, J., Truckenbrodt, H., and Jacobsen, T. (2012). The role of stimulus cross-splicing in an eventrelated potentials study. Misleading formant transitions hinder automatic phonological processing. The Journal of the Acoustical Society of America, 131(4):3120.

Strauss, M., Sitt, J. D., King, J.-R., Elbaz, M., Azizi, L., Buiatti, M., Naccache, L., van Wassenhove, V., and Dehaene, S. (2015). Disruption of hierarchical predictive coding during sleep. Proceedings of the National Academy of Sciences of the United States of America, 112(11):E1353–62.

Sun, Y., Giavazzi, M., Adda-Decker, M., Barbosa, L. S., Kouider, S., Bachoud-Lévi, A.-C., Jacquemot, C., and Peperkamp, S. (2015). Complex linguistic rules modulate early auditory brain responses. Brain and Language, 149:55–65.

Todorovic, A. and de Lange, F. P. (2012). Repetition Suppression and Expectation Suppression Are Dissociable in Time in Early Auditory Evoked Fields. Journal of Neuroscience, 32(39):13389–13395.

Traxler, M. J. (2014). Trends in syntactic parsing: Anticipation, Bayesian estimation, and good-enough parsing. Trends in Cognitive Sciences, 18(11):605–611.

Tyler, L. K. (1984). The structure of the initial cohort: Evidence from gating. Perception & Psychophysics, 36(5):417–427.

Van Petten, C., Coulson, S., Rubin, S., Plante, E., and Parks, M. (1999). Time course of word identification and semantic integration in spoken language. Journal of Experimental Psychology: Learning, Memory, and Cognition, 25(2):394–417.

Van Petten, C. and Luka, B. J. (2012). Prediction during language comprehension: Benefits, costs, and ERP components. International Journal of Psychophysiology, 83(2):176–190.

Wacongne, C., Changeux, J.-P., and Dehaene, S. (2012). A Neuronal Model of Predictive Coding Accounting for the Mismatch Negativity. Journal of Neuroscience, 32(11):3665–3678.

Wacongne, C., Labyt, E., van Wassenhove, V., Bekinschtein, T., Naccache, L., and Dehaene, S. (2011). Evidence for a hierarchy of predictions and prediction errors in human cortex. Proceedings of the National Academy of Sciences, 108(51):20754–20759.

Wilson, M. P. and Garnsey, S. M. (2009). Making simple sentences hard: Verb bias effects in simple direct object sentences. Journal of Memory and Language, 60(3):368–392.

Winkler, I. and Czigler, I. (2012). Evidence from auditory and visual event-related potential (ERP) studies of deviance detection (MMN and vMMN) linking predictive coding theories and perceptual object representations. International journal of psychophysiology : official journal of the International Organization of Psychophysiology, 83(2):132–43.

Winkler, I. and Schröger, E. (2015). Auditory perceptual objects as generative models: Setting the stage for communication by sound. Brain and Language, 148:1–22.

Yabe, H., Tervaniemi, M., Reinikainen, K., and Näätänen, R. (1997). Temporal window of integration revealed by MMN to sound omission. NeuroReport, 8(8):1971–1974.

Ylinen, S., Huuskonen, M., Mikkola, K., Saure, E., Sinkkonen, T., and Paavilainen, P. (2016). Predictive coding of phonological rules in auditory cortex: A mismatch negativity study. Brain and Language, 162:72–80.

